# A model of depth-dependent responses from neural superposition in fly compound eyes

**DOI:** 10.64898/2026.07.28.741303

**Authors:** Cora Hummert, Jouni Takalo, Vera Vasas, Mikko Juusola, Barbara Webb

**Author notes:** Current Address: School of Informatics, University of Edinburgh, Edinburgh, Scotland, UK.

## Abstract

Fly compound eyes pool signals from photoreceptors that sample the same region of visual space through neural superposition in the lamina. The optical axes of photoreceptors projecting to a single lamina cartridge are not perfectly parallel, but instead converge at a point a few millimeters in front of the eye. At short viewing distances (1–10 mm) this leads to distance-dependent differences in receptive field overlap. We explored whether it was possible that flies could sense depth in this “personal space” purely from the geometry of neural superposition. To this end, we combined a computational model of fly eye optics with a disparity-tuned lamina model originally developed for stereoscopic prey capture in praying mantises, and simulated the responses of lamina monopolar cells to moving stimuli at different distances. Across variations in stimulus parameters and lamina models, we found that lamina neuron responses indeed contain a distance-dependent component, which can overall be summarised as an enhanced response due to temporally overlapping receptor responses at a critical distance of 3–4 mm, the convergence distance of photoreceptor axes.

Depending on the lamina model and stimulus size, either response amplitude or onset gradient, or both, exhibited this peak. We further show that changes in eye size systematically shift this preferred distance, such that larger flies had a peak response at a greater distance. Our results demonstrate that lamina cell responses may contain a robust, geometry-derived component that is specific to object distance and invariant to other stimulus properties. This suggests that neural superposition, beyond improving sensitivity, may function analogously to a light-field camera system that is effectively “focused” on a behaviorally relevant distance.

**Author summary:** In this study, we explored how neural superposition in fly compound eyes may lead to an enhanced response to objects at a behaviorally relevant distance. The optical axis of neurally pooled photoreceptors, from neighbouring ommatidia, are not parallel, but converge at a point a few millimeters in front of the eye, which should lead to the strongest correlation of their signals at that distance. This idea was tested by combining a geometric model of the fly eye optics with a computational model of how the lamina cells process the responses of photoreceptors, and evaluating the output for a moving bright dot at different distances. We found that the simulated lamina cell has its fastest response, as measured by the onset gradient, to a stimulus at the distance where the photoreceptors converge (3–4 mm). This distance scales with the simulated eye size and corresponds to behaviorally relevant distances for fly behavior. This suggests that neural superposition may act to enhance the response to objects at a critical distance in the early visual processing of flies.

## Introduction

In dipteran compound eyes, multiple photoreceptors from different ommatidia project onto the same lamina cartridge, forming a neural superposition eye [1, 2]. This arrangement is classically assumed to pool signals from photoreceptors that sample the same point in visual space, thereby boosting sensitivity under dim conditions without losing spatial resolution [3]. However, the photoreceptor’s receptive fields do not perfectly overlap [4–6]. Measurements show that in dipteran flies the optical axes of photoreceptors wired to the same cartridge are not exactly parallel: they converge a few millimeters in front of the retina, with the exact distance varying between and within species [5, 7, 8]. Thus, the contributing photoreceptors do not sample identical visual directions. This raises the possibility that neural superposition may preserve information about object distance, at least over short, behaviorally relevant ranges.

### Lightfield sampling in compound Eyes

The optical arrangement of compound eyes can be compared to a lightfield (or panoptic) camera [9], which is often constructed by adding a microlens array in front of the sensor array [10] (See Bottom row of Fig. 1). To explain the key principle, consider first a standard single-lens imaging system, in which a set of light rays emerging in multiple directions from a single point in space are captured and refocused by the lens onto a single location on the image plane, providing a summed estimate of the total light intensity (Top left in Fig 1). Geometrically, the exact convergence of the light rays on the image sensor for an object at a given distance from a given lens depends on the focal length (distance from lens to image sensor). If these properties are mismatched, the rays will not exactly overlap on the image plane, producing blur (blur circles for different object distances areillustrated in Fig. 1). Consequently, it is possible to infer depth from focus, by finding, for a particular part of the scene, the focal length that provides the clearest focus, usually evaluated by maximum contrast [11, 12].

**Fig 1.**
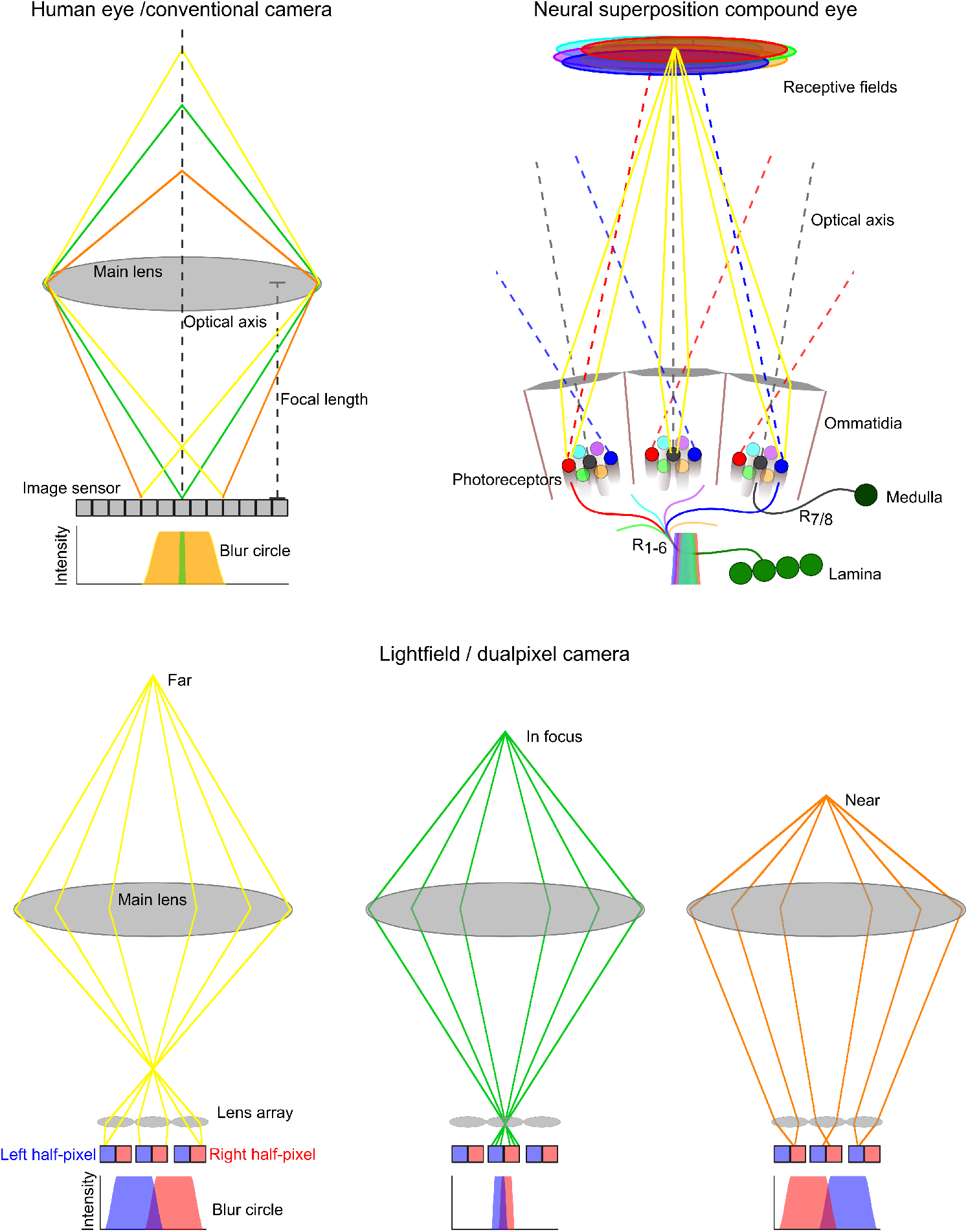
The optical set up for a conventional camera, a neural superposition eye and a dual pixel camera. The diagrams show light rays from a single point at different depths: far rays in yellow, in-focus rays in green and near rays in orange. Top left: conventional camera, the blur circle varies with depth but near and far blur circles are indistinguishable. Top right: Simplified, symmetric illustration of a neural superposition eye with perfectly overlapping receptive fields for the pooled photoreceptors *R*_1−6_ at a specific distance. The disparity of receptive field overlaps produces blur, which may preserve distance information for downstream visual circuits. Bottom: Lightfield / dual-pixel camera, the ’sorting’ of light rays preserves distance information, with the disparity of the blur circles on the left (Blue) and right (Red) half-pixel making objects equidistance near and far from the lens distinguishable.

In its most general form, a lightfield camera, instead of summing light rays from different directions, simultaneously or sequentially samples the light rays from specific directions, allowing retrospective reconstruction of the image corresponding to different focal lengths, and the extraction of depth (either by finding the focal length of maximum contrast as before, or by other methods exploiting the directional information). A simple example of this principle is dual-pixel focus found in many modern cameras. Each half of each pixel receives light rays from a different direction. Fig 1 shows how light rays emerging from different distances (left to right: far to near) are sorted by the microlens in front of the sensor array. If the scene is in focus, the input for each half-pixel will be the same; conversely objects not in focus produce a blur spread out over several half-pixels [13]. The spread of the blur circle on the half-pixels thereby provides depth information in a single exposure [14], as illustrated in the blur circles in the bottom row of Fig. 1.

In a similar manner, the individual photoreceptors of neighbouring ommatidia in the fly, which sample light rays from the same location but from different directions, could theoretically be used by downstream circuits to estimate depth. However, despite the apparent similarities to lightfield cameras, insect compound eyes encompass several different light processing principles. In the typical apposition eye, found in diurnally active insect such as beetles [15], there is minimal overlap in the field of view of the ommatidia, and a fused rhabdom conveys the summed light from all photoreceptors in one ommatidium to a corresponding lamina cell [16]. In the optic superposition eye, found for example in dragonflies [17], higher light sensitivity is obtained by an increased field of view, resulting in overlap, but optical and neural summing means that the directions of the incoming light rays is not preserved [18]. However, in neural superposition eyes (shown in Fig 1), overlapping fields of view project onto a small spatial array of receptors in each ommatidia, such that receptors from neighbouring ommatidia sample light rays from the same spatial region (or in other words, have overlapping receptive fields) but from different directions. In principle, this is lightfield sampling, and could be used to extract depth information, but the subsequent summation of these receptor outputs in the lamina cartridge suggests again that the relevant information is not preserved (see top right of Fig 1: Photoreceptor 1–6 project to one lamina cell).

The potential to extract depth from the differential light information received by overlapping receptive fields of the fly eye has been previously investigated in [7]. The key principle they use is to compare the outputs of pairs of receptors with parallel optical axes to pairs with converging or diverging optical axes, at a known angle, under the general assumption that a natural (static) scene contains approximately linear spatial intensity gradients. The difference in intensity detected by the parallel receptors provides a depth-independent measure of the gradient which can be used to normalise the depth-dependent difference between the non-parallel receptors, and thus recover a depth estimate. They use a geometric arrangement of receptors in a small array of ommatidia closely based on the eye of Musca domestica (see further below) and show that they can indeed produce depth estimates of images of objects in the range of 1.5–2 cm. However, this approach depends on comparing individual receptor outputs, which as noted above, does not seem consistent with the known neural circuits, which pool receptor responses.

### Distance tuning in near-field vision

We therefore consider the alternative possibility that the fly eye does not compute a general depth estimate, but instead exhibits a tuned response to a critical distance. To explore this idea, we adapt a model previously proposed to account for stereoscopic disparity-based distance estimation in mantises [19]. This model uses a disparity-tuned neuron that produces a maximal response for items of a specific size and distance, corresponding to prey that would evoke an attack strike by the mantis. In an analogous monocular model for the fly eye, the disparity tuning should leverage the lightfield sampling to produce an enhanced response at distances that are behaviorally relevant to the fly.

At long distances, angular offsets are negligible relative to the acceptance angle. At near distances, the offsets become large enough that each photoreceptor samples a measurably different region of visual space. Under bright-light adaptation, when acceptance angles shrink [4, 6, 20], these differences become even more pronounced. This results in a near range of 1–10 mm in which the overlap between photoreceptor receptive fields varies systematically with distance, and as a consequence, the joint photoreceptor output carries information about how far a visual object is from the eye.

This range corresponds to relevant behaviors. Fruit flies perform gap-crossing at 2–5 mm using monocular cues [21]. Courtship interactions also occur at 3–5 mm [22]. More generally, flies often interact with and manipulate objects with their forelegs [23], suggesting that determining if an object is ‘within reach’ might be ecologically important. Moreover, the personal space of a fly may differ both between and within species. For example, the gap crossing behaviour of flies is linked to their body size, with larger flies attempting crossing of larger gaps [24]. Larger flies also have larger compound eyes with smaller interommatidial angles, shifting the convergence point further away; smaller flies show the opposite pattern [8].

Here we explore the possibility that flies could sense depth in this near ‘personal space’ without binocular or motion parallax cues, but instead through the static geometry of neural superposition. We present a conceptual computational model showing that the pooled lamina response of a neural superposition eye can, in principle, provide distance-dependent information by producing enhanced responses to simulated moving objects at particular distances. We also simulate different stimulus sizes to test whether size has a comparable effect as in the stereoscopic prey capture model [19], but realistic stimulus sizes cover the whole visual field of the pooled receptors. The model therefore responds to the edge of the stimulus, and consequently the stimulus size effects observed in [19] are not expected to occur in our simulations.

We compare alternative models of lamina processing and also examine whether light-dependent (photomechanical) photoreceptor movements, driven by phototransduction reactions [25–27], which morphodynamically narrow and shift receptive fields [4, 6, 27–30], interfere with or enhance this superposition-based depth information. Finally, we show that increasing or decreasing eye size shifts the distance at which the enhanced response occurs, suggesting a simple geometric link between eye morphology and near-space visual tuning.

## Materials and methods

### Model Overview

We developed a computational model of a compound eye that combines the simplified fly optics as modeled in [7] (which in the following we refer to as the ‘*Bitsakos model* ’) with a disparity-tuned lamina cell model, in the following referred to as the ’*Adapted lamina model* ’, based on the model from [19]. This model follows the scheme of the stereoscopic disparity model in [19], but for neural superposition instead of binocular convergence. The distance estimation in [19] is based on a cell tuned to a critical binocular disparity, that is, receiving binocular input from two receptive fields that converge and overlap in a specific region of visual space. Instead of the binocular disparity, our adaptation of [19] uses the disparity between the receptive fields of individual photoreceptors projecting to one lamina neuron to model a depth-specific response. We include two variations of the model, the ‘*O’Keeffe model* ’, which conforms more closely to [19] and the ‘*Fu model* ’, which is an adaptation from the lamina cell modeled in [31].

The optics from the *Bitsakos model* represent the region of visual space from which each receptor collects light, including the overlap in receptive fields resulting from the specific optical axes and acceptance angles of the receptors in neighbouring ommatidia.

### Compound eye model

The *Bitsakos model* corresponds to [7] in the arrangement and response of photoreceptors and was implemented in Python 3.11. The model consists of a hexagonal ommatidial lattice with seven photoreceptors per ommatidium (*R*_1−7_, see top left of Fig. 2), each characterized by an optical acceptance angle and receptor-specific divergence.

**Fig 2.**
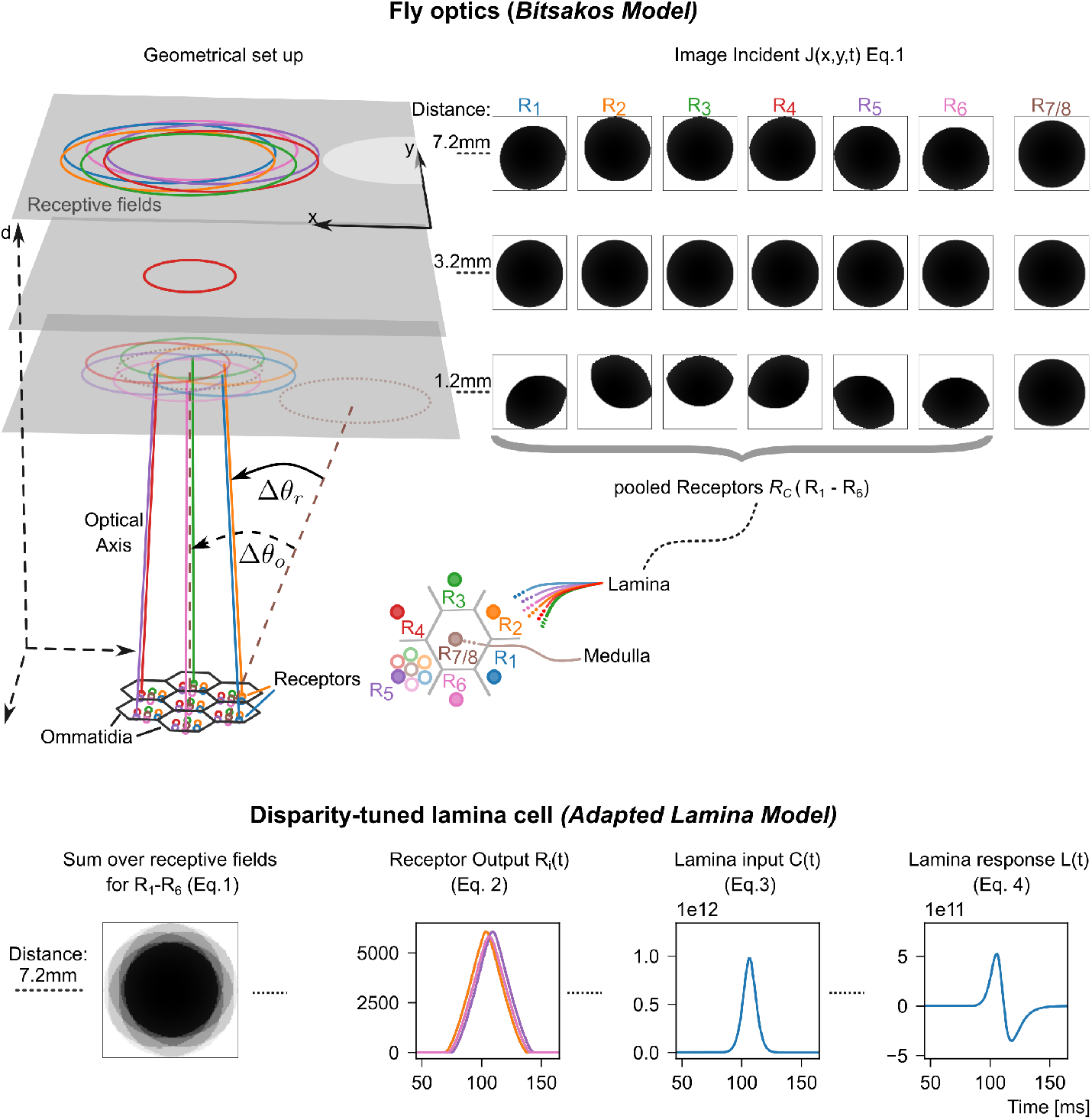
The model of the fly optics (*Bitsakos model*, top) and a disparity-tuned lamina cell (*Adapted lamina model*, bottom). The top left shows the geometrical set up of the simulated compound eye and stimulus on the screen, showing converging receptive fields from seven ommatidia (color coded). The optical axis of each ommatidia differs by the interommatidial angle Δ*θ_o_*and each receptor in an ommatidia has a divergence angle Δ*θ_r_*, as in [7]. The top right shows the overlap of these receptive fields with a small dot in the center of the screen at three distances (d=1.2, 3.2, 7.2 mm), which determines the input intensity to each receptor as calculated in equation 1. In the *Adapted lamina model* shown in the bottom, the responses of all photoreceptors projecting to a laminar cartridge are combined (left). The response over time to a moving dot for each receptor is the integral over its receptive field (column 2, equation 2) and the response of all receptors is summed compared to a threshold (0) and raised to a power (equation 3, column 3) providing the cartridge input. This is then highpass filtered to model the response of a lamina cell (equation 4, column 4).

We used a symmetrically arranged array of seven ommatidia (Top left of Fig. 2). Each ommatidium had a default diameter of 16 *µ*m, consistent with model parameters for a house fly used in [7], and contained seven receptors with one receptor (*R*_7_) at the center and the receptors one to six arranged symmetrically around the center (*R*_1−6_. Receptor divergence angles Δ*θ_r_* were defined as in [7], relative to the ommatidial axis, with interommatidial angle Δ*θ_o_*of 1.88^◦^ and each receptor had an acceptance angle Δ*θ_p_*of 1.9^◦^. In this arrangement the optical axis of the six photoreceptors *R*_1−6_ from the peripheral ommatidia that project to one lamina cartridge (see below) converge at approximately 3.2 mm in front of the eye (illustrated on the top left of Fig. 2.)

The spatial sensitivity of each receptor *R_i_*was modeled as a two-dimensional Gaussian kernel *f_G_*, with a width determined by the acceptance angle. The light intensities *J_i_*(*x, y, t*) seen by a photoreceptor *i* were computed as the convolution of this Gaussian Kernel with the presented stimulus *S*:

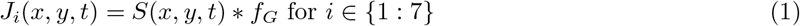

To normalise the total light input given the increasing size of the receptive field at increasing distances, the input *J_i_*(*x, y, t*) is sampled at 100 points. Example images for the light inputs of the six pooled photoreceptors are shown in Fig 2, top right. The integral over the receptive field (the sum over this sample) gives the response of each receptor *R_i_*over time.

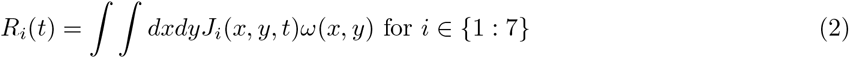

The response of the photoreceptors can be modulated by an additional spatial filter described by the weight matrix *ω*(*x, y*), for consistency with the model in [19]; but as explained below, we set these weights equal to one to model the photoreceptor response.

We also simulated a smaller and a larger eye by decreasing the ommatidia diameter *O_D_* and acceptance angle Δ*θ_p_*by 15% and increasing the interommatidial angle Δ*θ_o_* by 15% for a smaller eye and vice versa increasing/decreasing for a larger eye (*O_D_* = 0.016 mm ±0.0024, Δ*θ_p_* = 1.9^◦^ ± 0.28, Δ*θ_o_* = 1.88 ∓ 0.28). As the divergence angles are defined relative to the interommatidial angle in [7] and they need to be larger than the interommatidial angle for neural superposition to work, they were also increased by 15%. This variation of the interommatidial angle and ommatidia diameter builds on experimental results by [8] showing that eye size scales with fly body size (depending on feeding conditions around 12–24%) and that the number and size of ommatidia decreases with eye size while the interommatial angle increases, with the reverse being true for an increased eye size. In our model varying the ommatidial diameter and interommatidial angle accordingly leads to a scaled eyeradius *r* (*r* =0,3 mm for a smaller eye, *r* =0.5 mm for a larger eye, default *r*=0,4mm) and scaled distance at which the optical axis of the superimposed photoreceptors converge (approximately 2.3 mm for a smaller eye and 4.3 mm for a larger eye).

### Disparity-tuned lamina cell models

In [19], a binocular cell combines inputs from the two eyes with overlapping receptive fields at a specific disparity. The inputs are modeled as the result of several stages of visual processing preceding this convergence. The retinal image is obtained by convolving the visual input with a Gaussian with the standard deviation of the ommatidia acceptance angle, to mimic the resolution of an apposition eye. This retinal image is then highpass filtered to mimic phasic responses in the lamina. The input to the binocular cell is then calculated (for each eye) by convolving the image with a center–surround weighted receptive field *ω*(*x, y*), using an excitatory central region and an inhibitory surround, and summing over the receptive field.

In our *Adapted lamina model*, we simplified this processing as we are modeling the convergence that occurs already at the lamina in neural superposition compound eyes. As such, the ‘receptive field’ of each photoreceptor in our model is not centre-surround but a constant weight across the Gaussian, and the highpass filtering is assumed to occur after the input from the receptors has been combined in the lamina cartridge. An example for the receptor outputs and filtered response is shown in Fig 2.

Specifically, the instantaneous input *C*(*t*) to the lamina neuron *L* is the sum over all photoreceptors converging on a lamina cell (the peripheral photoreceptors of neighboring ommatidia *R_C_*, see Fig 1) minus tonic inhibition *b* and with exponent *γ* to control response selectivity.

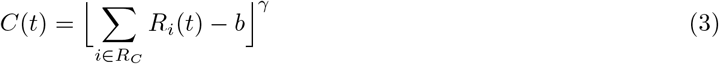

For parameters *b, γ* we used the same values as in [19]. The input is filtered with a first-order butterworth filter (implemented using the scipys lfilter function) with a lower cutoff frequency of *l* = 4*Hz* and an upper cutoff frequency of *h* = 60*Hz*. The cutoff frequency was chosen to be significantly slower than the sampling rate and we varied the cutoff frequencies of the butterworth filter to match the lamina response from [19].

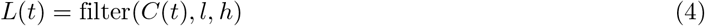

To assess how modeling the lamina cell and cartridge affects the response to stimulus distance, we also tested two variations to this model: The *O’Keeffe model* is more consistent to [19], i.e., with a center-surround receptive field and temporal filtering introduced for individual photoreceptors before summing in the lamina cartridge (See Supporting information for an illustration of the processing). The parameters of the receptive field in this model are adjusted from [19] to have a larger excitatory region (Width inner excitatory region = 30 samples, Width outer excitatory region = 60 samples). Otherwise parameters are the same as in [19]. Secondly, we tested a different lamina cell model, the *Fu model* based on [31], in which the temporal filtering for the lamina cell is done as follows, with the delay *n_p_* = 20:

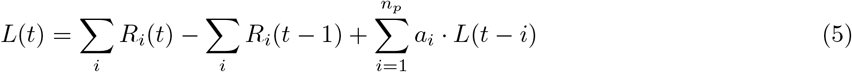

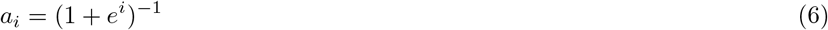

We only model the first layer of the lamina cells from [31] to test how a different temporal filter affects the disparity tuning. [31] modeled the visual processing of the fly up to the lobula plate, but disregard the optics of neural superposition in their model of receptor inputs. For a final test of the sensitivity of the *Adapted lamina model* to parameter changes in the pooling, we modeled the cartridge as a sum without inhibition, thus simplifying Eq. 3 further to

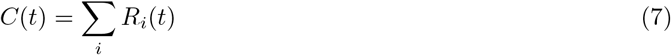

### Photoreceptor movement

Our initial analysis uses static photoreceptors (which in the following we refer to as ’non-moving Rc’), but motivated by the photoreceptor model of *Drosophila melanogaster* in [4] we also explored the effects of light-induced photoreceptor movement (in the following refered to as ’moving Rc’). In response to light input the photoreceptors have been observed to move inside the ommatidia on a path away from the lens, narrowing the receptive field, and sideways, shifting the receptive field sideways. The photoreceptors in the left eye move to the right, thereby shifting the receptive field to the left and vice versa in the right eye. The receptive field motion thus follows the optic flow for a forward moving fly. In [4] this was modeled as a spring-damper system with the activation *act*(*t*) driving the movement according to the sum over the absorbed photons of the seven photoreceptors of one rhabdomere:

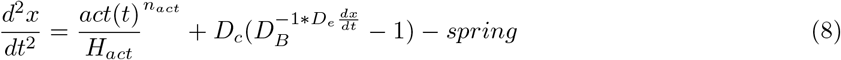

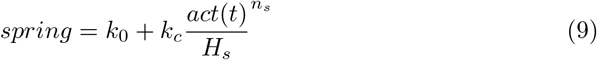

We use the the same equation for the movement and model the activation as the sum over the light intensities *J_i_*_=1:7_(*x, y, t*) seen by the photoreceptors of one rhabdomere *Rhb* on one ommatidium.

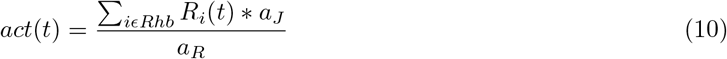

Since the model in [4] had maximal photon flux of 9000 *ph/µm*1/2 and the modeled movement became unstable for higher inputs, the activation in our implementation was normalized to lie below 350 with *a_J_* = ∑*_iɛRhb_ J_i_*_=1:7_(*x, y*) for *J_i_*_=1:7_(*x, y*) = *f_G_*(a stimulus intensity of one covering the entire receptive fields). All other parameters are the same as in [4] (supp. Material, page 87). As our model is symmetrical with the same acceptance angles for all receptors, this photoreceptor movement moves and narrows the receptive fields by the same amount for all receptors, which corresponds to a shift along the x-axis on the screen by approximately half the width of a receptive field.

### Stimulus Generation

Visual stimuli were presented on a virtual screen located at variable distances (1.2–7.2*mm*) in front of the eye. Thus any one stimulus was presented at one distance from the eye. The screen had a simulated pixel size of 0.4*µm* and was updated with a temporal resolution of 1 ms for all simulations. The tested stimulus was a bright dot (Stimulus intensity = 1.0 on a screen intensity = 0.0) moving parallel to the compound eye. We tested a dot of a size of 90% (0.9 va or 1.7^◦^) and 390% (3.9 va or 7.4^◦^) of the receptive field size, moving at 50, 75 and 100^◦^*/s*. The dot size was set in percentage of the receptive field size, so that for every distance the image of the stimulus on the receptor would be the same. This was done to test whether the difference in overlap could be used to disambiguate larger objects at a farther distance from near, smaller objects. In the range of the simulated distances the larger dot size might be considered to correspond to realistic object sizes, with a size of 1 mm at a distance of 10 mm. The smaller dot size was only a few *µ*m in size and mainly simulated to test the principle of a distance specific response in Lamina cells. We also varied the direction of the stimulus (+1, i.e., front-to-back or −1, back-to-front) and the movement axis (horizontal (x) or vertical (y), see top screen of Fig. 1 for an illustration of the stimulus movements). When included, the receptor movement was modeled as a positive shift of the receptive field on the screen; the receptive fields thus moved with the stimulus for a stimulus moving along the x-axis in direction +1 (back-to-front).

### Distance Analysis

To measure the response of the modeled Lamina cell to a stimulus at a specific distance, we integrate the rectified *L*(*t*) over the duration of the stimulus, as also done in [19].

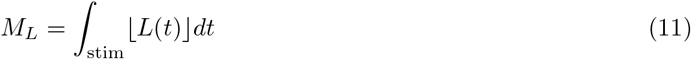

As another measure, we calculated the mean temporal derivative of the lamina response during the first 10 ms (*N* = 10) following the first positive derivative of the response (in the following referred to as the onset gradient *G*_onset_). This measure represents how quickly a downstream neuron might reach the threshold to fire an action potential and is defined as follows with *t*_+_ as the first time point at which the temporal derivative of the lamina response *L*(*t*) became positive.

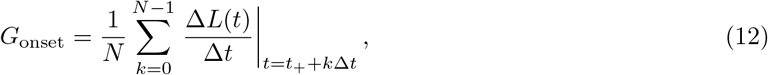

## Results

We simulated a neural superposition eye using the *Bitsakos model* of the compound eye [7] and combined it with a disparity-tuned lamina cell model derived from a stereoscopic prey capture model [19] to investigate the distance dependence of neural superposition. The left panel of Fig 3 shows, in 2D, the optical geometry for two photoreceptors (R3 and R6, see Fig. 2 for photoreceptor arrangement) that project to the same neural cartridge, with their separation and optical axes as defined in the methods, but with the acceptance angle Δ*θ_p_* set to 0.4^◦^ to make the visualization of the region of overlap more salient. The area in which the receptive fields of the two photoreceptors overlap is indicated by the distances *d*_far_ (the furthest distance at which overlap starts) and *d*_near_ (the nearest distance where overlap ends). The relationship between the acceptance angle and these two distances is shown in the second panel: for acceptance angles larger than 0.6^◦^, *d*_far_ is infinite, i.e. the receptive fields continue to overlap at all distances larger than *d*_near_. This holds for the acceptance angle used in our model (third panel) but the degree of overlap still varies with distance, and falls off sharply for smaller distances (right panel in Fig 3). Irrespective of the acceptance angle, the receptive fields overlap perfectly at a distance of 3.2mm. This distance-dependent change in the receptive field overlap should result in differences in the temporal alignment of the photoreceptor outputs for a moving stimulus but also a difference in contrast of still images at different distances.

**Fig 3.**
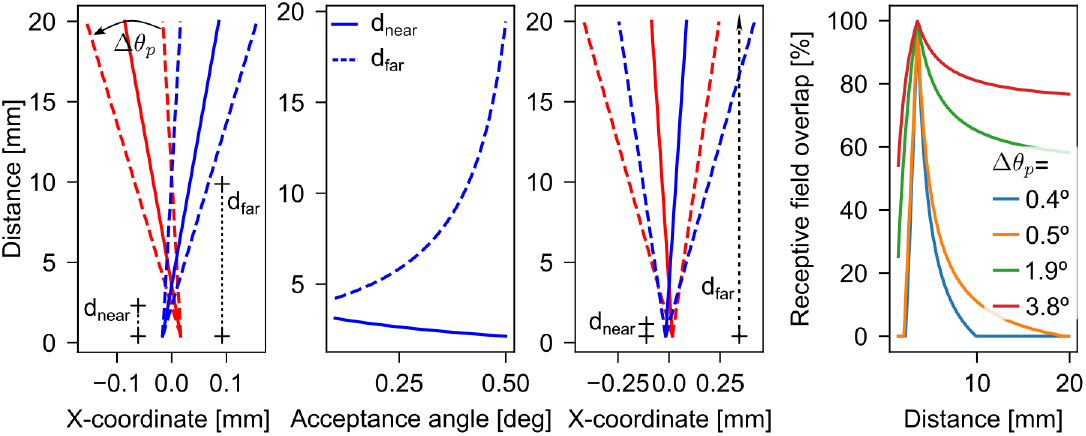
How receptive field overlap varies with distance. For simplicity we show the geometrical relation for two receptors from neighbouring ommatidia with overlapping receptive fields, illustrated in 2D. Left: The optical axis (solid line) and the receptive field size for an acceptance angle Δ*θ_p_* = 0.4 for two receptors positioned on the zero x-axis (R6: blue, R3: red). The maximal distance at which the receptive fields overlap (*d_far_*) and the minimal distance (*d_near_*) of overlap are marked. Middle left: The distances *d_far_* (dashed), *d_near_* (solid), are plotted against the acceptance angle Δ*θ_p_*. Middle right: The geometrical set up for an acceptance angle of Δ*θ_p_* = 1.9 as used in the current model (as illustrated in leftmost panel). The receptive fields always overlap for distances higher than 0.7 mm, therefore *d_far_* is infinite. Right: The overlap of the receptive field of two receptors as a percentage of the whole visual field seen by the two receptors, plotted against distance, for four acceptance angles.

### Effect of stimulus distance on lamina response

To test whether neural superposition can in principle generate distance-dependent signals, we first examined the *Adapted lamina model* ’s response for non-moving photoreceptors and a stimulus smaller than the receptive fields. Within a range of distances *<* 8*mm* we simulated a bright dot of size 0.9 visual angle moving parallel to the photoreceptors, which would correspond to an object only a few *µ*m in size. In Fig 4 we visualise the response of all the receptors in one cartridge (left column) for stimuli at increasing distances (rows). As the degree of receptive field overlap changes, the degree of blur of the stimulus changes, with the sharpest edge at distance 3.2 mm (Column 1, Row 3 in Fig 4). This is also visible in the timecourse of the photoreceptor outputs: At distances greater or smaller than 3.2 mm, some receptor outputs (e.g. *R*_4_ and *R*_5_, with *R*_5_ plotted on top of *R_d_*in Column 2 Fig 4) already begin to decrease before all receptors (e.g. *R*_1_ and *R*_2_) have reached their maximal activation, resulting in a broader summed receptor response and smaller lamina input at stimulus onset. At the convergence distance, the receptor outputs peak simultaneously, producing the strongest lamina input (3.2 mm, Row 3 in Column 4 Fig 4).

**Fig 4.**
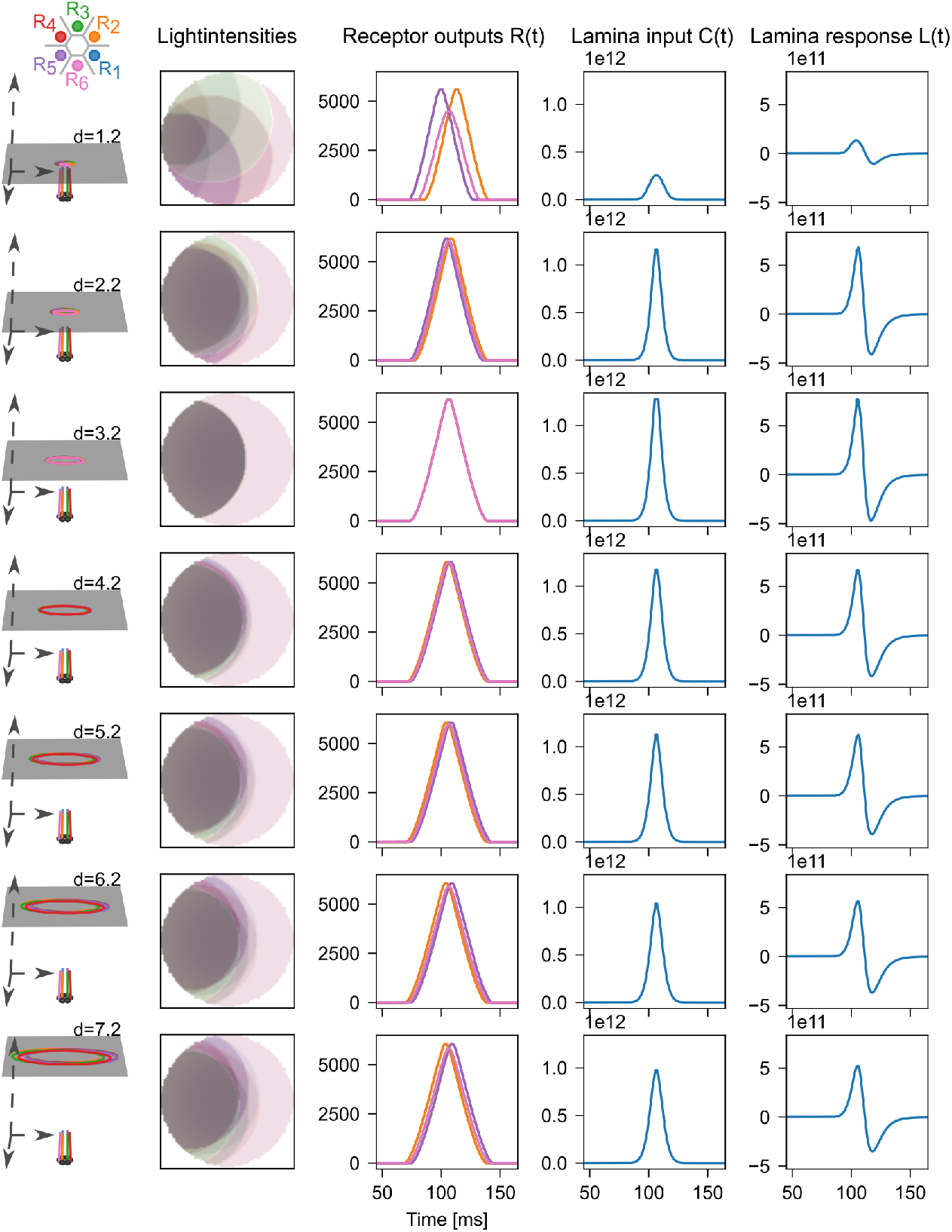
Depth-dependent responses in the model for a small moving stimulus. The model outputs for the receptors of one neural cartridge for a circle moving across a screen at different distances (rows, set up illustrated in the first column) to the eye. The stimulus moves along the positive x-axis, front-to-back with a speed of 50^◦^*/s* (stimulus size= 0.9 va). The second column shows the overlaid images of the six pooled receptive fields as the stimulus enters. Due to the symmetric arrangement of the receptors and Ommatidia (receptor positions in the eye shown on the top left), receptor outputs are plotted on top of each other with only R2, R5 and R6 visible. The third column shows the receptor outputs over time, the fourth shows the lamina input and the fifth column the lamina response for the *Adapted lamina model*. The temporal dispersion of photoreceptor responses is smallest for 3.2mm, resulting in the highest lamina response at this distance.

The change in response with distance is also visible when integrating over the rectified lamina response, shown in Fig 5: The lamina cell response is maximal for 3.2 mm. We also calculated the gradient of the Lamina response and took the mean of the gradient over the first 10 ms of a positive response. As in the integrated Lamina response, the onset gradient (defined in Eq. 12) is highest for the distance at which the converging photoreceptors perfectly overlap (bottom row of Fig 5. While the stimulus direction had no effect on the lamina response (Lines for the two stimulus directions in Fig. 5 are plotted on top of each other), the peak of the integrated lamina response and gradient is slightly higher for a stimulus moving along the y-axis. This is due to the orientation of the ommatidia, which are arranged symmetrically around the center ommatidia, but therefore not symmetrically aligned with the cardinal directions of the stimulus movement (see Fig 2). A smaller stimulus moving along the y-axis thus moves right through the center of the receptive fields, while a stimulus moving along the x-axis passes through slightly off center.

**Fig 5.**
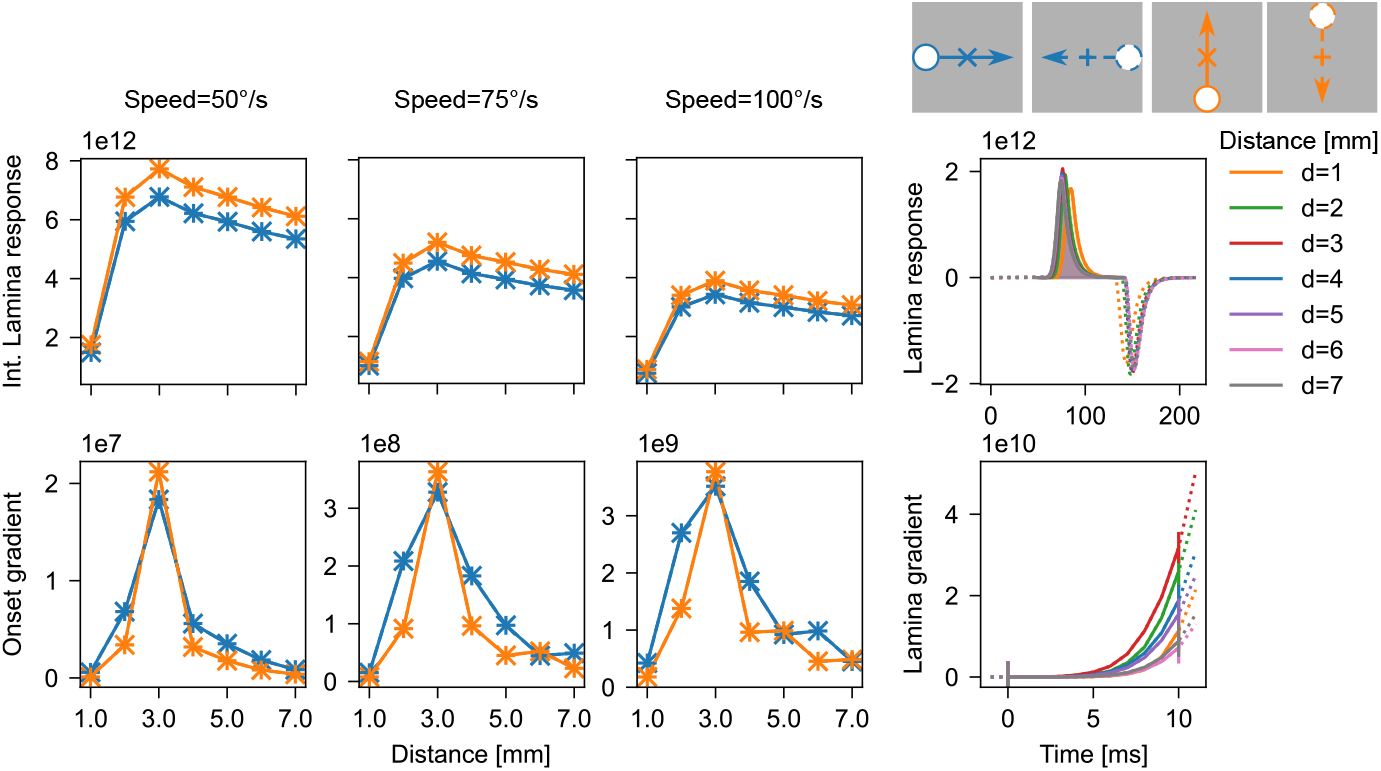
The lamina response is maximal at a specific distance. The rectified response integrated across the stimulus duration *M_L_* (top row, calculation illustrated on the right) and the onset gradient *G*_onset_ (bottom row, calculation illustrated on the right) of the modeled lamina cell in the *Adapted lamina model* for different screen distances. Both *M_L_* and *G*_onset_ peak at the convergence distance (3.2 *mm*). The stimulus is a small circle (size: 0.9 va) moving at three speeds (50 − 100^◦^*/s*, columns) along the four cardinal directions on the screen (See illustration at the top: Color indicates the movement axis of the stimulus and linestyle the stimulus direction). Due to the symmetric set-up the response is the same for both stimulus directions and lines are plotted on top of each other.

### Testing larger stimuli and alternative lamina cell models

A more realistic stimulus size for objects seen by the fly (e.g. the head of another fly) is approximately 1mm, which corresponds to 3.9 va at 10 mm in the model. As shown in the first column in Fig 6, the edge of the larger stimulus is blurred for distances smaller and larger than 3.2 mm, similar to the smaller stimulus.

**Fig 6.**
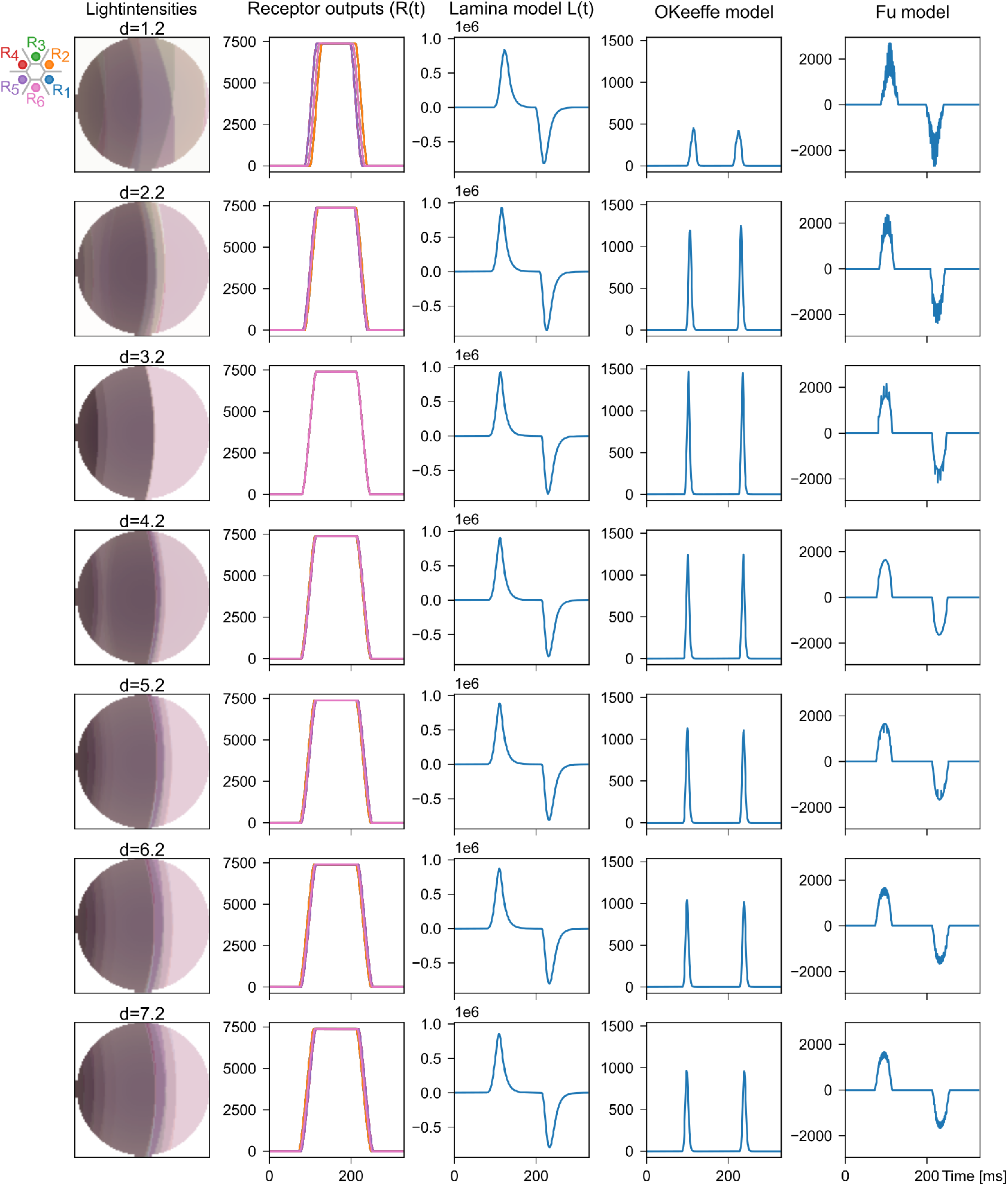
Alternative lamina models also show depth-dependence. The outputs of the three different models for a dot (size=3.9va, overlaied image shown in first column) moving front-to-back at 50^◦^*/s* along the x-axis of a screen at different distances (distances vary with rows). The second column shows the non-moving Receptor outputs projecting to one Cartridge for all distances, with receptor outputs plotted on top of each other and only receptors R2, R5 und R6 visible due to the symmetric arrangement of the receptors. The third, fourth and fifth column show the Lamina cell response as modeled with the *Adapted lamina model*, the *O’Keeffe model* and the *Fu model*. The temporal dispersion at stimulus onset results in a maximal response for the convergence distance in the *O’Keeffe model*, but only weak modulation of the lamina response peak in the *Adapted lamina model* and the *Fu model*

However, unlike the smaller stimulus, the larger stimulus covers all receptive fields simultaneously for most of its trajectory.

For the larger stimulus, the onset of the receptor responses remains distance dependent, with receptor activation spread out over time for distances larger or smaller than 3.2 mm (Column 2, Fig. 6). However, once the stimulus fully covers the receptive fields, all receptors reach maximal activation regardless of distance. The summed receptor input therefore differs primarily during stimulus onset and offset, while the central part of the response becomes nearly identical across distances. Consequently, distance-dependent effects are weaker in measures based on the total lamina response and are expressed more strongly in measures that quantify the initial response dynamics.

### Lamina cell models

All model variants hyperpolarized in response to a positive light increment, but differed in the temporal profile of their responses (Fig. 6). Of the three models, the *Adapted lamina model* most closely resembles experimentally measured lamina cell responses [32]. The *Fu model* shows a slower return from hyperpolarization, whereas the inhibitory surround in the *O’Keeffe model* produces a small depolarization at stimulus onset. In addition, the squaring operation in the *O’Keeffe model* removes the depolarization observed in lamina cells in response to light decrements.

These differences also affect how strongly distance-dependent changes are expressed. In our Adapted model, receptor outputs are summed before temporal filtering, so the largely overlapping receptor responses produce only small differences in the lamina response across distances (Column 3, Fig. 6). The *Fu model* behaves similarly, although its temporal filter amplifies small fluctuations in the summed receptor signal more strongly. In contrast, the *O’Keeffe model* applies temporal filtering before summation. As a result, receptor responses are dominated by stimulus onset and offset, making the summed receptor activity and lamina response visibly dependent on stimulus distance (Column 4, Fig. 6).

To quantify these effects, we compared the integrated positive lamina response (Eq. 11) and the onset gradient (mean gradient over the first 10 ms of a positive lamina gradient; Eq. 12. Illustrated on the right in Fig. 5). The *O’Keeffe model* showed a clear peak in the integrated response at the convergence distance of 3.2 mm for all stimulus directions and sizes as well as a peak at the convergence distance for the onset gradient.

In the *Fu model*, the integrated response peaks at 3.2 mm for the small stimulus but becomes nearly distance-independent for the larger stimulus. The onset gradient, however, remains maximal at the convergence distance (Middle row of Fig. 7). The same pattern is observed in the *Adapted lamina model*, where the onset gradient consistently peaks at 3.2 mm despite only small differences in the integrated response (Bottom row of Fig. 7). The simplified *Adapted lamina model* that excluded the tonic inhibition in Eq. 3 showed the same behaviour as the *Adapted lamina model*, with a peak at the convergence distance for the onset gradient for both stimulus sizes and no strong distance modulation for the integrated lamina response for the large stimulus (See supplementary material, Fig **??**).

**Fig 7.**
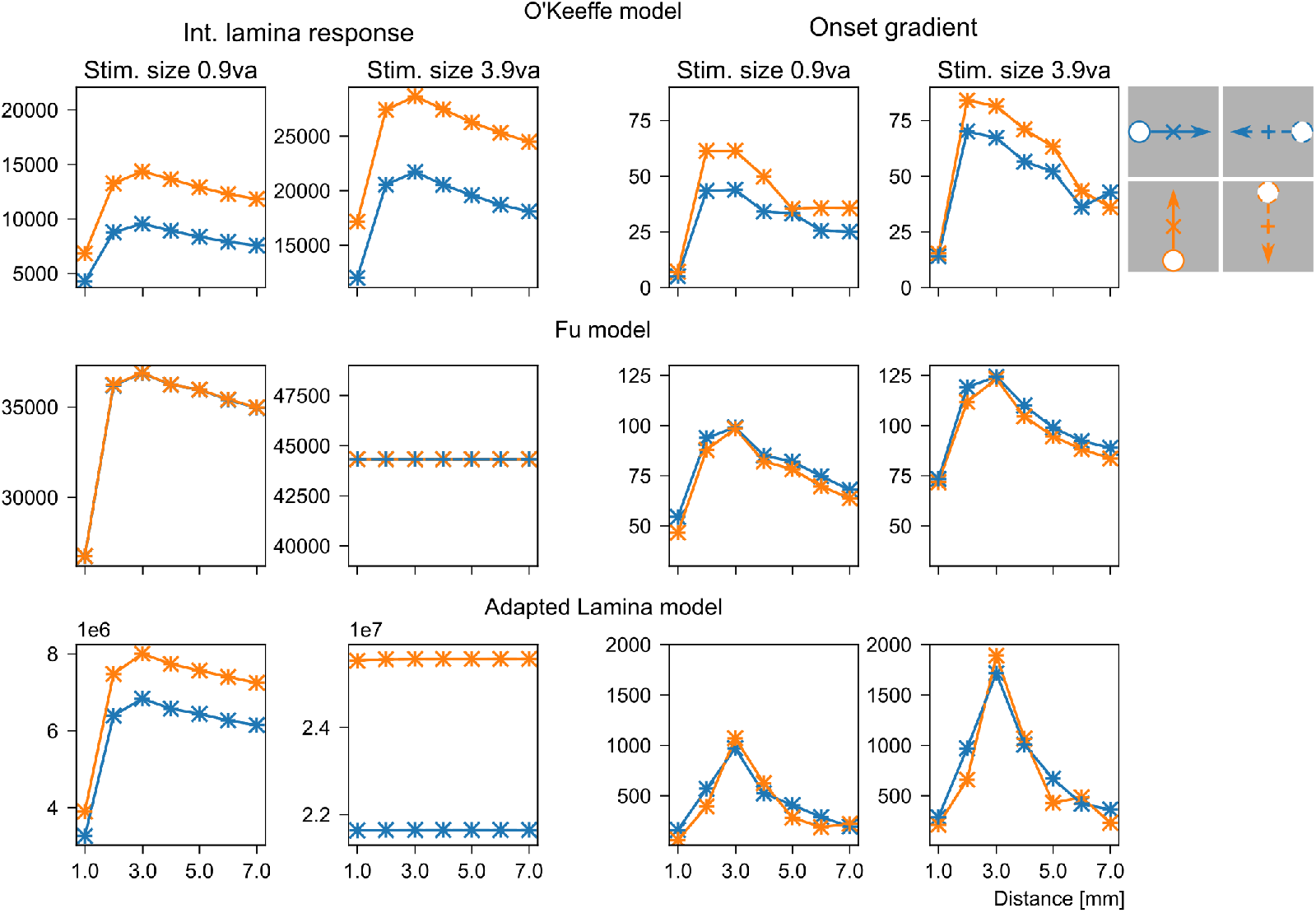
Distance response for model variants. The integrated lamina response *M_L_*and mean of the lamina gradient *G*_onset_ is plotted against the simulated screen distance for different lamina cell models (top row: *O’Keeffe model*, middle row: *Fu model*, bottom row: *Adapted lamina model*). The stimulus speed is 50^◦^*/s* and stimulus movement axis and direction are indicated by linecolor and linestyle respectively, with the lines for the stimulus direction plotted on top of each other. The long period of full overlap of receptor responses for the larger stimulus (see column 2, Fig. 6) overwhelms any distance dependence in the integrated laminar response for the *Fu model* and *Adapted lamina model*. The onset gradient peaks at the convergence distance for all sizes and model variations.

Overall, all models exhibited a distance-dependent effect in our onset gradient measure. The differences in the integrated lamina response for model variations can be explained by the temporal filtering properties of the respective models and by whether filtering occurs before or after summation of the photoreceptor signals.

### Effects of photoreceptor movement

The larger stimulus covers the whole receptive field of all photoreceptors of one rhabdomer as it passes the center of the receptive fields. All receptors of one ommatidia will thus be activated roughly at the same time, which leads to movement of the rhabdomer. We therefore compare the moving and non-moving photoreceptors in our *Adapted lamina model* for the larger stimulus.

In our *Adapted lamina model* with the photoreceptor movement included, the receptor outputs also vary with stimulus direction. For stimulus moving back-to-front and against the photoreceptor movement, the receptor and lamina response is shorter and has a steeper gradient in comparison to the stimuli moving with the receptive fields (see Fig 8). This change of the receptor outputs is consistent with [4], who found a prolonged response for stimuli moving front-to-back and thus with receptive field motion and optic flow and a shorter and steeper response for stimuli moving against receptor motion. Additionally, we observe double-peaked responses for stimuli moving with the receptive fields, which makes it difficult to extract a single measure for distance tuning.

**Fig 8.**
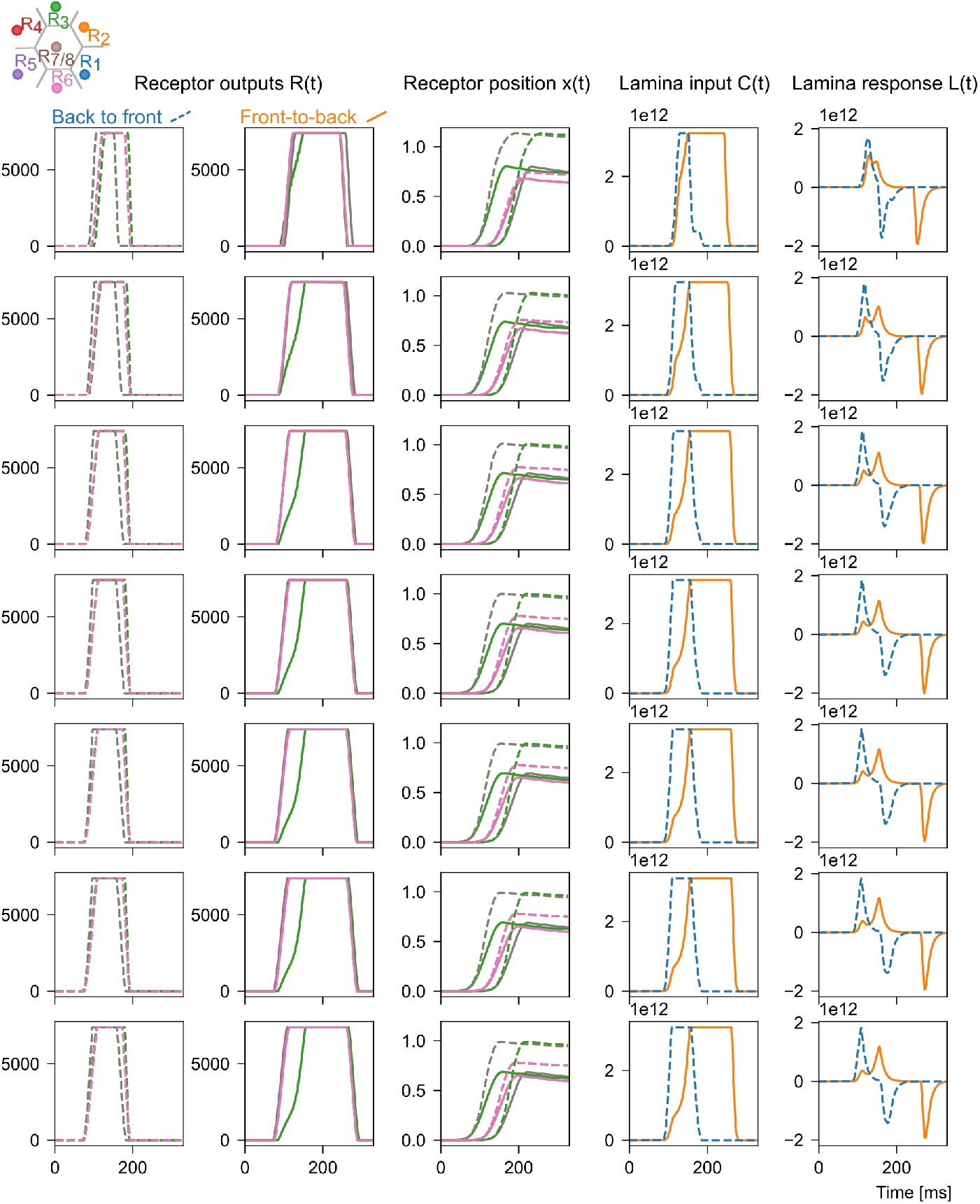
Moving receptors change the laminar response. The outputs of the moving receptors for a dot (size=3.9 va, speed=50^◦^*/sec*) moving back-to-front (dashed line, first column) and front-to-back (solid line, second column) along the x-axis on a screen simulated at different distances (Rows: Distances 1.2–7.2 mm). The first and second column show the receptor outputs for the pooled receptors and receptor *R*_7*/*8_ (Gray) for both stimulus directions and the third column show their positions over time. The forth and fifth column show the lamina input and response as calculated in the *Adapted lamina model* in Eq. 4 and Eq. 3 respectively. The receptor movement results in a prolonged receptor output and lamina response for stimuli moving with the receptive fields (front-to-back) and a shorter, steeper response for stimuli moving against stimulus direction (back-to-front).

Nevertheless, the onset gradient for moving photoreceptors and a stimulus moving along the x-axis remained maximal at distances between approximately 2 and 4 mm, with a direction- and speed-dependent modulation (Fig. 9). For stimuli moving against receptor motion the onset gradient has significantly higher peaks. Because receptor movement changes the profile of the lamina response, including the appearance of double-peaked responses, the shifts of the peaks may reflect limitations of the metric rather than meaningful shifts in preferred distance. Importantly, receptor movement did not abolish the distance-dependent effect. For stimuli moving along the y-axis, where receptor movement should have little influence, the onset gradient continued to peak at the same distance as in the non-moving receptor simulations (bottom row in Fig. 9).

**Fig 9.**
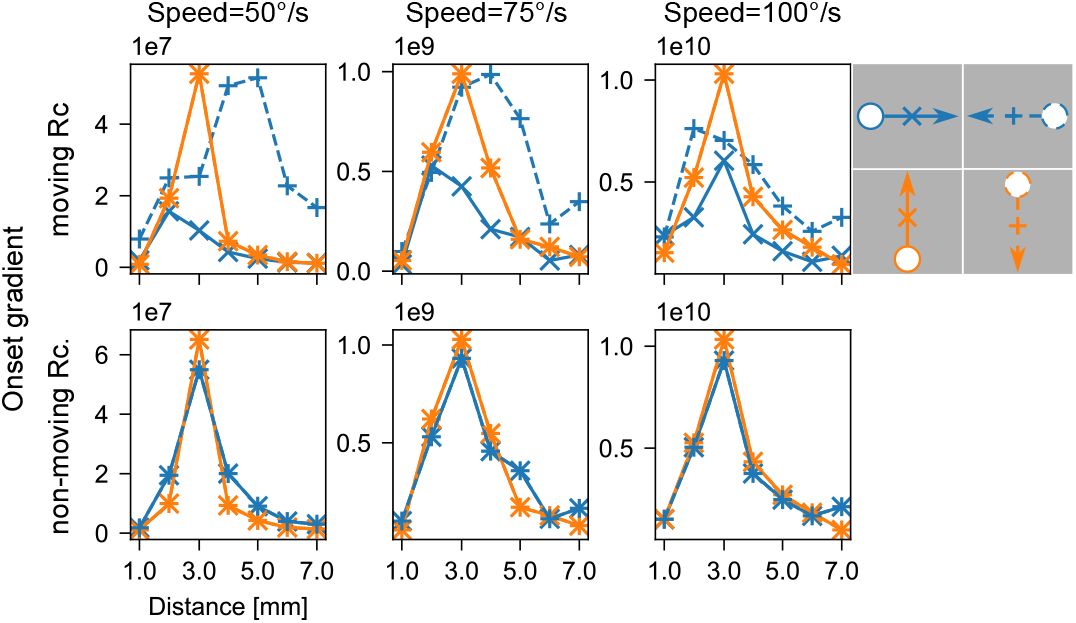
Moving receptors still produce distance-dependent lamina onset gradients. The onset gradient for different stimulus speeds (50 − 100^◦^*/s*, columns) for the *Adapted lamina model* with moving receptors, plotted against the simulated screen distance. The onset gradient peaks at distances ranging from 2.2-4.2 mm for moving Receptors and at 3.2 mm for non-moving Receptors. Stimulus movement axis and direction are indicated by linecolor and linestyle respectively, with lines for the two stimulus directions plotted on top of each other.

### Juusola Model

Juusola, Song and Takalo have developed biophysically realistic, morphodynamic, multiscale models of quantal sampling and processing in fly compound eyes, based on systematic information-theoretic analyses of experimental data [4, 6, 27, 28, 33–37]. Their latest models for Drosophila melanogaster photoreceptors [4] use experimentally estimated receptor positions and divergence angles for each photoreceptor and are thus not symmetric, but physiologically more accurate. This means that the receptive fields of pooled photoreceptors never perfectly overlap. The model also includes the photoreceptor movement (as modeled in Eq. 9), the mechanisms for photon absorbtion, a stochastic quantum bump model and the Hodgkin-Huxley membrane model for the voltage response of the receptors. To test whether a more physiological realistic model may also show a distance specific response for the pooled photoreceptors, we simulated a bright dot, smaller than the receptive field size at four distances (1.25, 2.5, 5, 10mm) and moving along the cardinal directions. We used the simulated receptor voltages of one neural cartridge as an input to our *Adapted lamina model*. As the photoreceptor voltages are negative, the model parameters are adapted to that input as follows: The threshold to in Eq. 4 is set to *b* = −380, *γ* = 3. The temporal filtering is done with matlabs Butterworth filter function *filter*. The photoreceptor voltages shown in the left column of Fig 10 are spread out over roughly 80ms for all distances. While the temporal dispersion of different photoreceptors varies with distance, there is no obvious systematic distance-dependent trend. However, the lamina response differs slightly but visibly with distance (top right panel Fig 10), with the smallest peak for a distance of 1.25mm. Consistent with these results, the integrated lamina response increases sharply between 1.25 and 2.5 mm but then continues to increase gradually or level out for longer distances. The onset gradient predominantly varies with the movement axis and direction of the stimulus, but also shows a distance dependent component with a peak at 5 mm for a stimulus moving up and thus against receptive field motion. Thus, although the distance dependence is substantially weaker than in the symmetric model, a measurable modulation of the lamina response with distance remains present in the physiologically more realistic simulation which includes photoreceptor movement.

**Fig 10.**
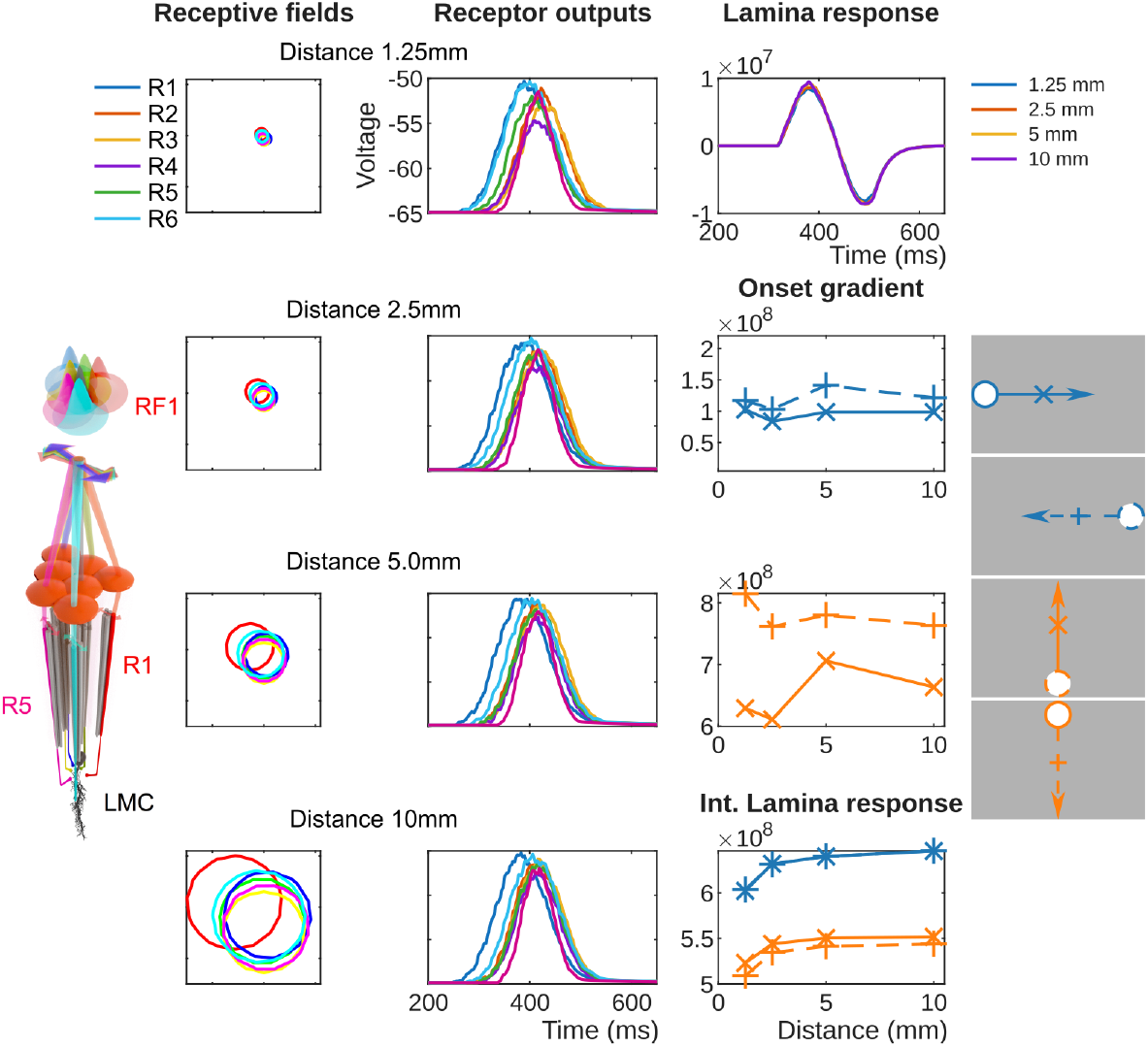
Physiologically exact photoreceptor model. The Lamina response for a physiologically exact Photoreceptor model [4] for a small moving dot (speed=50^◦^*/s*, size=3va) at different distances from the eye. The Rhaddomere with the photoreceptors and receptive fields is illustrated on the left. The first column shows the receptive fields on a screen at different distances (1.25–10.0 mm) and the second column the respective photoreceptor voltages for the dot moving with the receptive field. The top right panel shows the corresponding lamina response for all distances (color coded). The two middle panels on the right show the onset gradient for all cardinal directions (Top: Stimulus movement along x-axis, Bottom: Stimulus movement along y-axis and the bottom right panel shows the integrated Lamina response. Stimulus movement axis and direction are also indicated by linecolor and linestyle as in figure 9). The receptive field overlaps and photoreceptor voltages show small variations with distance and the integrated lamina response increases with distance. The onset gradient varies with distance, with a peak for a stimulus moving against photoreceptor movement along the y-axis.

### Variations of Eye Size

Finally, we tested how eye size influences distance tuning by simulating a smaller (ommatidia diameter *d* = 0.013 mm (−15%)) and larger (ommatidia diameter *d* = 0.018 mm (+15%)) compound eyes, following the morphological variation reported by Currea [8]. Since the photoreceptors were arranged symmetrically around the center of the ommatidia, the optical axis of pooled photoreceptors all converged at one point, (2.3 mm for a smaller eye and 4.3 mm for a larger eye). As we were mainly interested in the effect of the eye size on the distance tuning, we simulated only one stimulus speed (75^◦^*/s*) and movement (along y-axis, back-to-front) for each size. The peak of the onset gradient was shifted with the eye size and decreased for larger eyes with a larger acceptance angle (see Fig 11). The smaller gradient can be explained by the resampling of the receptive fields: In the simulations with a larger acceptance angle the stimulus moves through a larger region, thus taking longer, but the receptive field outputs are resampled to have the same sample size of 100x100.

**Fig 11.**
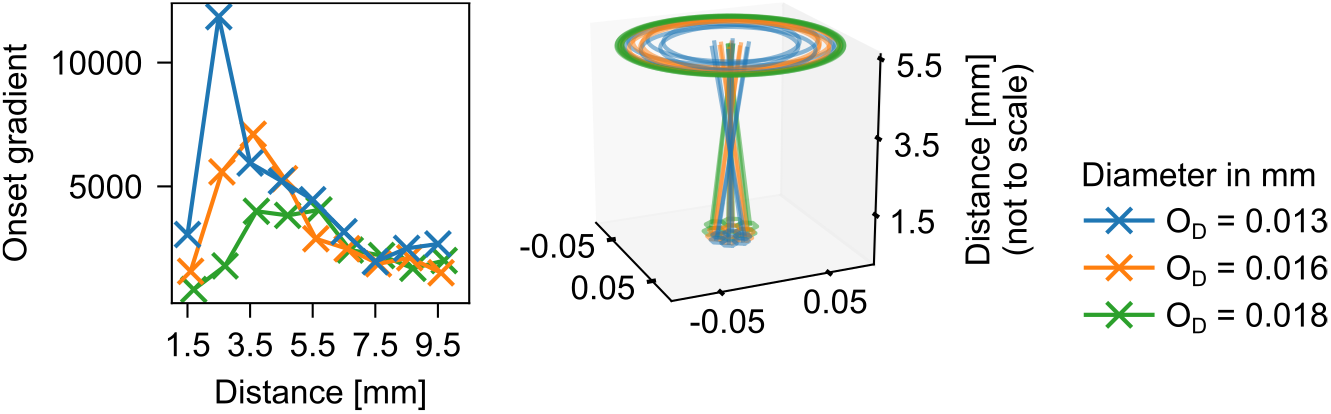
Peak response distance changes with eye size. The onset gradient of the *Adapted lamina model* for different simulated eyes sizes (color coded, Blue *O_D_* = 0.013 mm, Orange *O_D_* = 0.016 mm, Green *O_D_* = 0.018 mm) plotted against the simulated screen distance. The changes in the geometry of the eye as modeled by increasing the ommatidia diameter *O_D_* and decreasing interommatidial angle shift the convergence distance, as illustrated on the right. The peaks of the onset gradient are shifted with the simulated eye size. The stimulus is moving back-to-front along the y-axis of the screen.

## Discussion

We developed a simple model that combined the geometry of the neural superposition compound eye in *Musca* [7] with processing to detect when a visual object is a specific distance from the eye, based on a model of mantis prey detection [19]. We found that a distance-specific response of the simulated lamina cell persists across model variations and different stimuli parameters. Most notably, in our *Adapted lamina model* the mean gradient at onset of the modeled lamina response (onset gradient) was maximal near the distance at which the optical axes of the pooled photoreceptors converged. This distance scaled systematically with eye size and corresponds to the distance at which flies interact with objects, courtship song is performed and gaps are crossed (around 1.7*x* of body size) [22, 23, 38].

### Distance-dependent responses across lamina cell models

We implemented several variants of the lamina cell model that differed in how temporal filtering was applied relative to the pooling of photoreceptor signals. Despite these differences, all models exhibited a distance-dependent response component, indicating that the effect arises from the geometry of neural superposition rather than from a specific implementation of lamina processing.

The strongest distance dependence was observed in the *O’Keeffe model* [19], in which the individual photoreceptor signals are temporally filtered before being summed. Both the integrated lamina response and the onset gradient therefore peaked at the convergence distance of the pooled photoreceptors. However, this sequence is not physiologically realistic. In neural superposition eyes, photoreceptor signals from different ommatidia first converge onto the same lamina cartridge, where synaptic transmission and network processing then shape the pooled response. The pooling of photoreceptors is not expected to be a simple linear summation and responses in the lamina exhibit phasic nonlinear interactions and gain-control mechanisms [39, 40, 40–42]. These could enhance the distance-dependence of the lamina response in a similar manner to the *O’Keeffe model*. Consistent with this possibility, synaptic high-frequency jumping demonstrates that lamina processing does not simply average photoreceptor inputs, but nonlinearly transforms them into fast, behaviourally relevant temporal signals with exceptionally high information content [6]. Distance-related optical structure may likewise be dynamically reshaped and amplified rather than averaged away.

Our *Adapted lamina model* and the *Fu model* [31] are closer to the dipteran visual circuity because photoreceptor signals are first pooled and subsequently filtered. In both models, distance-dependent effects were weaker in the integrated lamina response, but the onset gradient consistently peaked near the convergence distance. This modulation of the gradient suggests not so much a stronger but a faster response to objects at a certain distance.

Although the actual circuitry of the fly lamina is considerably more complex than the simplified models used here, the persistence of the effect across multiple model variants suggests that distance-dependent modulation is a robust consequence of the underlying optical geometry itself. Taken together, these results indicate that neural superposition naturally generates distance-dependent temporal structure in the pooled photoreceptor signal, which could, in principle, be available to downstream visual circuits.

### Photoreceptor movement

We also explored the effects of light-induced photoreceptor microsaccades [4, 6, 27, 28, 30], both in our *Adapted lamina model* and in a more physiologically detailed model [4]. The receptor movements introduced strong direction-dependent modulation of the lamina response for stimuli moving parallel to the direction of receptor displacement. Nevertheless, the distance-dependent component in the onset gradient remained present, which suggests that the receptor motion does not break neural superposition. While the increase of the onset gradient with stimulus speed and for stimuli moving against the receptor movement is consistent with [4, 6, 28], the observed shifts in preferred distance remain difficult to interpret as a biologically meaningful interaction between receptor motion and receptive field overlap. Our chosen metrics may thus be able to capture a distance-specific effect for moving receptors while being too limited for a complete analysis.

As the interaction between receptor microsaccades and neural superposition remains difficult to interpret within the present simplified framework, we also tested a more physiologically more realistic model that simulated photoreceptor responses [4]. In addition to the miccrosaccades, the optical axes in this model converge at slightly different distances around 3–4 mm, which means the receptive fields never perfectly overlap. Given this, it is not surprising the distance-dependent effect is substantially weaker and mostly appears between the smallest and farthest distance. That a small distance-dependent modulation of the lamina response remained, suggests that neither photoreceptor motion nor a not perfect receptive-field convergence completely eliminate the distance dependence of neural superposition.

### Implications for fly behaviour

The observed distance-dependence in our model emerged directly from the geometry of the eye and consequently shifted with morphological variations. For smaller eyes with a smaller ommatidial diameter and larger ommatidial angle [8], the optical axes of pooled photoreceptors converged at nearer distances, while the convergence distance of larger eyes is farther away. The onset gradient of the lamina responses shifted accordingly with peaks at distances proportional to eye size. This scaling is interesting because eye size varies naturally with body size in flies [8] and body size correlates distances relevant to fly behaviour, such as gap crossing [24].

Gap crossing in flies occurs at distances of approximately 2–5 mm and relies primarily on monocular visual cues [21, 38]. Courtship interactions also occur within a similar spatial range [22]. More generally, flies frequently interact with nearby objects using their forelegs [23], making near-field object localization behaviourally important. Rather than explicitly encoding depth, neural superposition may therefore bias visual processing toward objects located within this ‘personal space’. In this interpretation, nearby objects would evoke stronger or faster responses simply because their image geometry better matches the convergence properties of the eye. Rather than explicitly computing object distance, the early visual system would be most sensitive to objects located within a characteristic near-field region determined by the geometry of neural superposition.

### Neural superposition as a biological light-field sensor

An alternative interpretation of these results is that neural superposition produces a distance-dependent focus effect. In our symmetric model, the pooled image formed by photoreceptors projecting to a single cartridge is sharpest at the convergence distance of the optical axes and becomes progressively blurred at nearer or farther distances. Although perfect overlap disappears in more realistic eye geometries, the same principle remains: there exists a range of distances for which the pooled image is most ’focussed’.

In principle, neural superposition therefore behaves similarly to a focus-tuned optical system, producing maximal contrast and a faster response for objects located within a specific range of distances. A similar principle to neural superposition has been applied to lightfield cameras in [10], where the information of overlapping images from the lightfield camera was fused to obtain a high-resolution image. While in the fly eye the information from individual receptors is lost in the neural superposition, distance-specific information that is only available because of the lightfield sampling by the photoreceptors is preserved.

The distance-tuned response we observed is therefore a specific consequence of how neural superposition eyes pool lightfield information and not a general property of superposition eyes. While optical superposition eyes may also exploit their own lightfield sampling capacity the mechanism for this would have to be different.

### Limitations

Several limitations of the present study should be noted. The optical geometry in our main model was simplified and based largely on symmetric receptor arrangements derived from [7]. Real fly eyes exhibit asymmetries in receptor divergence angles and acceptance angles, as demonstrated by the more physiologically realistic models [4, 6]. Likewise, the lamina circuitry was represented only by simplified temporal filtering rather than detailed cellular models. Our simulations also used highly controlled stimuli rather than naturalistic visual scenes. Finally, although we demonstrate that distance-dependent response structure emerges naturally from neural superposition, we do not identify downstream circuits capable of decoding or exploiting this information behaviourally.

## Conclusion

Despite these limitations, the present work suggests a reinterpretation of neural superposition. Rather than simply increasing sensitivity by pooling redundant inputs, neural superposition may precisely preserve the small directional differences required to selectively enhance responses to nearby objects. In this sense, the fly eye may function analogously to a low-resolution biological light-field sensor, in which anatomy and early pooling circuitry together generate distance-dependent temporal structure in visual responses. Such a mechanism would provide a simple and computationally inexpensive way of emphasizing objects within a behaviorally relevant near-field space.

## Supporting information

**Fig 12.**
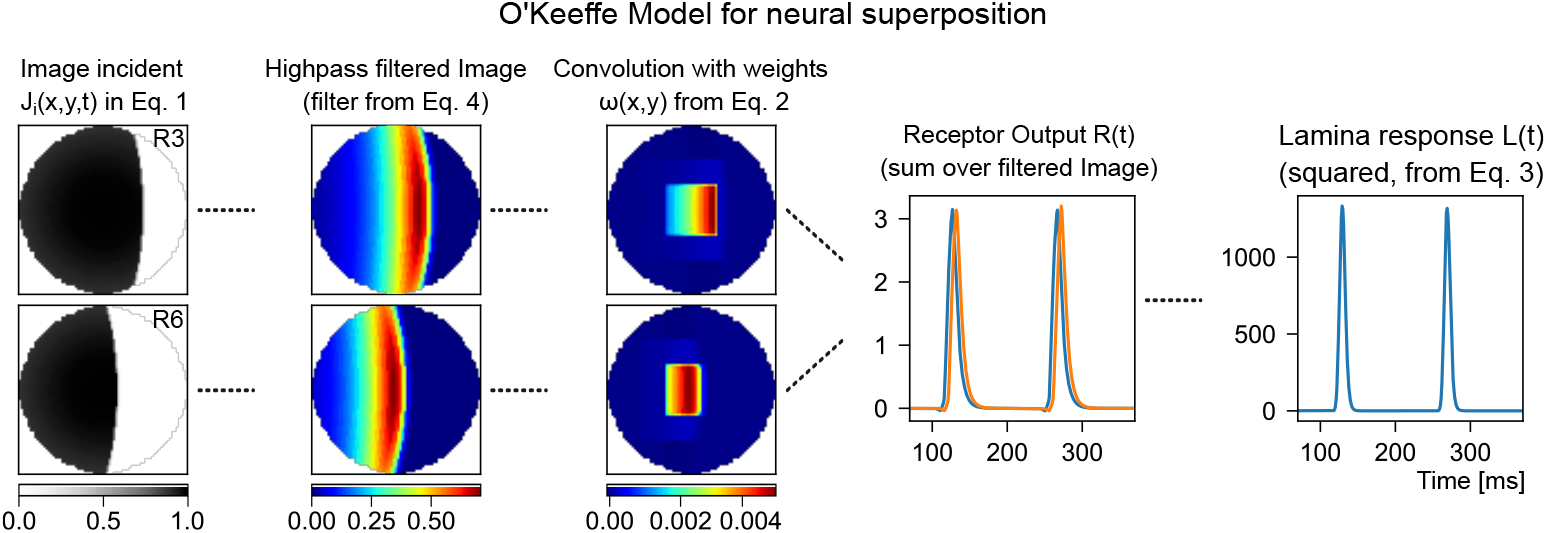
Illustration of the *O’Keeffe model* for neural superposition. The light intensities are highpass filtered as described in equation 4, shown in the second column, convolved with an excitatory-center-inhibitory-surround field (column three) and summed (equation 2 for *J_i_*(*x, y, t*) being the filtered images, shown in column four). The instantaneous response, called *v_R/L_* in O’Keeffe and *R*(*t*) here is modulated with a tonic inhibition and raised to a power (column five, equation 3).

**Fig 13.**
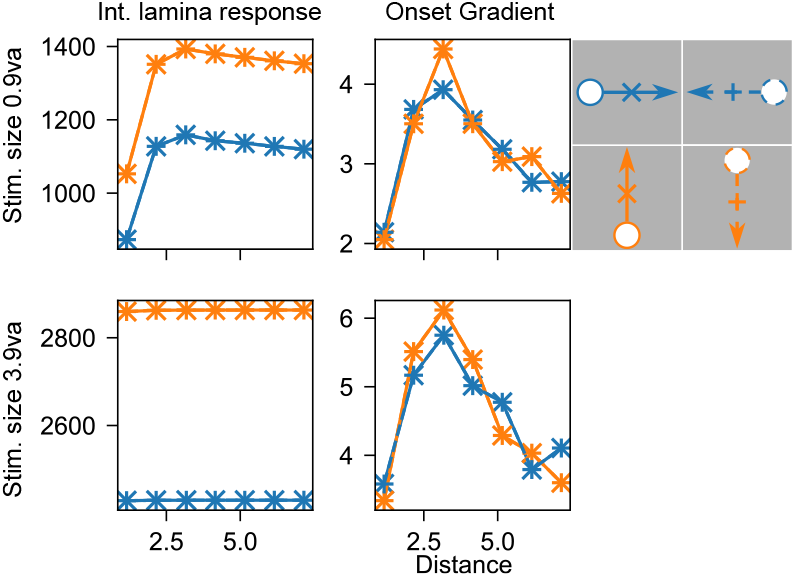
Distance response for the simplified *Adapted lamina model*. The integrated lamina response (1st column) and onset gradient (2nd column) plotted against the simulated screen distance for the *Adapted lamina model* with Eq 3 simplified to *C*(*t*) = ∑*_i_ R_i_*(*t*) (summed receptor response). Stimulus movement axis and direction are indicated by linecolor and linestyle respectively (lines for stimulus direction plotted on top of each other) and stimulus size varies with rows. The modulation of both the onset gradient and the integrated lamina response by the stimulus distance match the modulation of the *Adapted lamina model*, with the onset gradient peaking at the convergence distance.

## References

1. Kirschfeld K. [Neural superposition eye]. Fortschritte der Zoologie. 1973 1;21:229–57. Available from: https://europepmc.org/article/med/4762854.

2. Stavenga DG. The neural superposition eye and its optical demands. Journal of Comparative Physiology A. 1975 12;102:297–304. doi:10.1007/BF01464342.

3. Greiner B. Adaptations for Nocturnal Vision in Insect Apposition Eyes. International Review of Cytology. 2006 1;250:1–46. doi:10.1016/S0074-7696(06)50001-4.

4. Kemppainen J, Scales B, Haghighi KR, Takalo J, Mansour N, McManus J, et al. Binocular mirror–symmetric microsaccadic sampling enables Drosophila hyperacute 3D vision. Proceedings of the National Academy of Sciences of the United States of America. 2022 3;119:e2109717119. doi:10.1073/pnas.2109717119.

5. Pick B. Specific misalignments of rhabdomere visual axes in the neural superposition eye of dipteran flies. Biological Cybernetics. 1977 12;26:215–24. doi:10.1007/BF00366593.

6. Mansour N, Takalo J, Kemppainen J, Bridges AD, MaBouDi H, Bohra AA, et al. Synaptic high-frequency jumping synchronises vision to high-speed behaviour. Nature Communications 2026 17:1. 2026 5;17:3863. doi:10.1038/s41467-026-72509-2.

7. Bitsakos K, Fermüller C. Depth estimation using the compound eye of dipteran flies. Biol Cybern. 2006;95:487–501. doi:10.1007/s00422-006-0097-1.

8. Currea JP, Smith JL, Theobald JC. Small fruit flies sacrifice temporal acuity to maintain contrast sensitivity. Vision Research. 2018 8;149:1–8. doi:10.1016/J.VISRES.2018.05.007.

9. Feng X, Ma Y, Gao L. Compact light field photography towards versatile three-dimensional vision. Nature Communications. 2022 6;13:3333. doi:10.1038/s41467-022-31087-9.

10. Bishop TE, Favaro P. The light field camera: Extended depth of field, aliasing, and superresolution. IEEE Transactions on Pattern Analysis and Machine Intelligence. 2012;34:972–86. doi:10.1109/TPAMI.2011.168.

11. Zhuo S, Sim T. Defocus map estimation from a single image. Pattern Recognition. 2011;44(9):1852–8.

12. Favaro P, Soatto S. A geometric approach to shape from defocus. IEEE Transactions on Pattern Analysis and Machine Intelligence. 2005;27(3):406–17.

13. Wadhwa N, Garg R, Jacobs DE, Feldman BE, Kanazawa N, Carroll R, et al. Synthetic depth-of-field with a single-camera mobile phone. ACM Transactions on Graphics. 2018 7;37. Available from: https://dl.acm.org/doi/pdf/10.1145/3197517.3201329.

14. Kim D, Jang H, Kim I, Kim MH. Spatio-focal bidirectional disparity estimation from a dual-pixel image. In: Proceedings of the IEEE/CVF Conference on Computer Vision and Pattern Recognition; 2023. p. 5023–32.

15. Meyer-Rochow VB. Compound eyes of insects and crustaceans: Some examples that show there is still a lot of work left to be done. Insect Science. 2015;22(3):461–81. doi:10.1111/1744-7917.12117.

16. Land M, Nilsson D, Land M, Nilsson D. Apposition compound eyes. Animal Eyes. 2012:157–90.

17. Nilsson DE. The diversity of eyes and vision. Annual Review of Vision Science. 2021;7(1):19–41.

18. Land MF. Microlens arrays in the animal kingdom. Pure and Applied Optics: Journal of the European Optical Society Part A. 1997 nov;6(6):599. doi:10.1088/0963-9659/6/6/002.

19. OKeeffe J, Yap SH, Llamas-Cornejo I, Nityananda V, Read JCA. A computational model of stereoscopic prey capture in praying mantises. PLOS Computational Biology. 2022 5;18:e1009666. doi:10.1371/JOURNAL.PCBI.1009666.

20. Stavenga DG. Angular and spectral sensitivity of fly photoreceptors. III. Dependence on the pupil mechanism in the blowfly Calliphora. Journal of Comparative Physiology A. 2004 1;190:115–29. doi:10.1007/s00359-003-0477-0.

21. Pick S, Strauss R. Goal-driven behavioral adaptations in gap-climbing Drosophila. Current Biology. 2005 8;15:1473–8. doi:10.1016/j.cub.2005.07.022.

22. Coen P, Clemens J, Weinstein AJ, Pacheco DA, Deng Y, Murthy M. Dynamic sensory cues shape song structure in Drosophila. Nature. 2014 3;507:233–7. doi:10.1038/nature13131.

23. Durrieu M, Dall’osto D, Ka T, Lam C, Lobato-Rios V, Ramdya P. Object manipulation and affordance learning in Drosophila. bioRxiv. 2026 5:2026.04.28.721021. doi:10.64898/2026.04.28.721021.

24. Krause T, Spindler L, Poeck B, Strauss R. Drosophila Acquires a Long-Lasting Body-Size Memory from Visual Feedback. Current Biology. 2019 6;29:1833–41.e3. doi:10.1016/J.CUB.2019.04.037.

25. Hardie RC, Franze K. Photomechanical responses in Drosophila photoreceptors. Science. 2012 10;338:260–3.

26. Hardie RC, Juusola M. Phototransduction in Drosophila. Current Opinion in Neurobiology. 2015 10;34:37–45. doi:10.1016/J.CONB.2015.01.008.

27. Juusola M, Dau A, Song Z, Solanki N, Rien D, Jaciuch D, et al. Microsaccadic sampling of moving image information provides Drosophila hyperacute vision. eLife. 2017 9;6. doi:10.7554/ELIFE.26117.

28. Kemppainen J, Mansour N, Takalo J, Juusola M. High-speed imaging of light-induced photoreceptor microsaccades in compound eyes. Communications Biology 2022 5:1. 2022 3;5:203. doi:10.1038/s42003-022-03142-0.

29. Juusola M, Takalo J, Kemppainen J, MaBouDi H, Bu BY, Shukla S, et al. Beyond Static Perception: Animals, Neurons and Synapses Move to Compute Efficiently. 2026 3. Available from: https://www.preprints.org/manuscript/202603.1093. doi:10.20944/preprints202603.1093.v1.

30. Juusola M, Takalo J, Kemppainen J, Haghighi KR, Scales B, McManus J, et al. Theory of morphodynamic information processing: Linking sensing to behaviour. Vision Research. 2025;227:108537.

31. Fu Q, Yue S. Modelling Drosophila motion vision pathways for decoding the direction of translating objects against cluttered moving backgrounds. Biological cybernetics. 2020 10;114:443–60. doi:10.1007/s00422-020-00841-x.

32. Freifeld L, Clark DA, Schnitzer MJ, Horowitz MA, Clandinin TR. GABAergic Lateral Interactions Tune the Early Stages of Visual Processing in Drosophila. Neuron. 2013 6;78:1075–89. doi:10.1016/J.NEURON.2013.04.024.

33. Juusola M, Polaviejaz GGD. The Rate of Information Transfer of Naturalistic Stimulation by Graded Potentials. Journal of General Physiology. 2003 8;122:191–206. doi:10.1085/JGP.200308824.

34. Juusola M, Hardie RC. Light Adaptation in Drosophila PhotoreceptorsI. Response Dynamics and Signaling Efficiency at 25°C. Journal of General Physiology. 2001 1;117:3–25. doi:10.1085/JGP.117.1.3.

35. Juusola M, Hardie RC. Light Adaptation in Drosophila Photoreceptors: II. Rising Temperature Increases the Bandwidth of Reliable Signaling. Journal of General Physiology. 2000 12;117(1):27–42. doi:10.1085/jgp.117.1.27.

36. Song Z, Juusola M. Refractory sampling links efficiency and costs of sensory encoding to stimulus statistics. Journal of Neuroscience. 2014;34(21):7216–37. Available from: https://www.jneurosci.org/content/34/21/7216.short.

37. Song Z, Postma M, Billings SA, Coca D, Hardie RC, Juusola M. Stochastic, adaptive sampling of information by microvilli in fly photoreceptors. Current Biology. 2012 8;22:1371–80. doi:10.1016/j.cub.2012.05.047.

38. Triphan T, Nern A, Roberts SF, Korff W, Naiman DQ, Strauss R. A screen for constituents of motor control and decision making in Drosophila reveals visual distance-estimation neurons. Scientific Reports. 2016 6;6:27000. doi:10.1038/srep27000.

39. Laughlin SB, Howard J, Blakeslee B. Synaptic limitations to contrast coding in the retina of the blowfly Calliphora. Proceedings of the Royal Society of London B Biological Sciences. 1987 9;231:437–67. doi:10.1098/RSPB.1987.0054.

40. van Hateren JH. Neural superposition and oscillations in the eye of the blowfly. Journal of Comparative Physiology A. 1987 11;161:849–55. doi:10.1007/BF00610226.

41. Juusola M, Weckstrom M, Uusitalo RO, Korenberg MJ, French AS. Nonlinear models of the first synapse in the light-adapted fly retina. Journal of Neurophysiology. 1995;74:2538–47. doi:10.1152/JN.1995.74.6.2538.

42. Zheng L, Polavieja GGD, Wolfram V, Asyali MH, Hardie RC, Juusola M. Feedback Network Controls Photoreceptor Output at the Layer of First Visual Synapses in Drosophila. Journal of General Physiology. 2006 5;127:495–510. doi:10.1085/JGP.200509470.

