## Supplementary figures and images for "A model of depth-dependent responses from neural superposition in fly compound eyes"

### Supplemental Figure 1

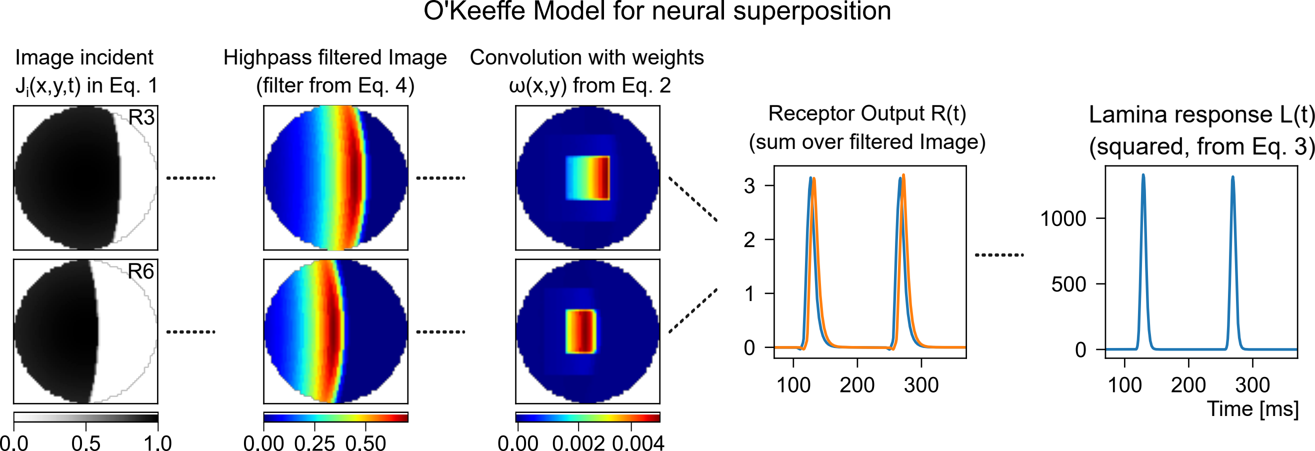

### Supplemental Figure 2

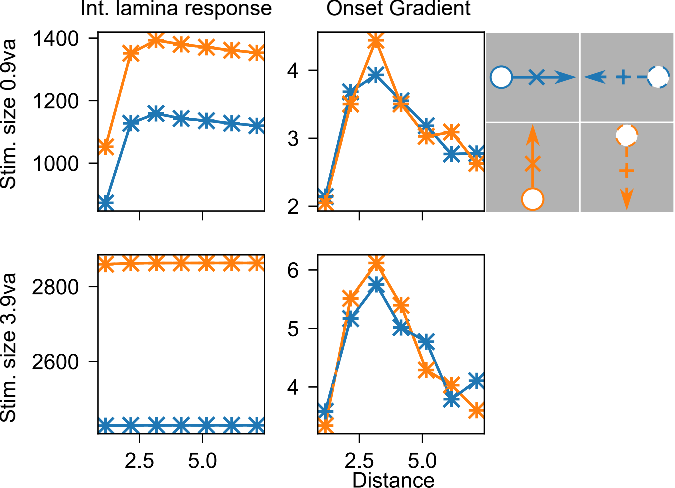
